# SARS-CoV-2 genome sequences from late April in Stockholm, Sweden reveal a novel mutation in the spike protein

**DOI:** 10.1101/2020.08.03.233866

**Authors:** Tatiany Aparecida Teixeira Soratto, Hamid Darban, Annelie Bjerkner, Maarten Coorens, Jan Albert, Tobias Allander, Björn Andersson

## Abstract

Large research efforts are going into characterizing, mapping the spread, and studying the biology and clinical features of the severe acute respiratory syndrome coronavirus 2 (SARS-CoV-2). Here, we report four complete SARS-CoV-2 genome sequences obtained from patients confirmed to have the disease in Stockholm, Sweden, in late April. A variant at position 23463 was found for the first time in one genome. It changes an arginine (R) residue to histidine (H) at position 364 in the S1 subunit of the spike protein. The genomes belonged to two different genetic groups, previously reported as two of the three main genetic groups found in Sweden. Three of them are from group B.1/G, corresponding to the Italian outbreak, reported by the Public Health Agency of Sweden to have declined in prevalence by late April, and more investigation is needed in order to ensure that the spread of different types of SARS-CoV-2 is fully characterized.

**Highlights:**

- Four near-complete genomes of SARS-CoV-2 were assembled from late April in Stockholm.
- A novel mutation in the spike protein were found.
- The phylogeny of the strains were discussed.

---

A severe pneumonia disease 2019 (COVID-19) caused by severe acute respiratory syndrome coronavirus 2 (SARS-CoV-2), emerged in Wuhan, China, in December 2019 and has rapidly spread around the world (Wu et al., 2020; Zhou et al., 2020). In Sweden, at the time of writing, more than 78,504 cases have been reported and 5,667 deaths have been confirmed (FHM, 2020). As this virus is new to humans, large research efforts are going into characterizing the virus, mapping its spread, and studying its biology and clinical features. We here report four complete SARS-CoV-2 genome sequences obtained from patients confirmed to have the disease. The sampling and tests were carried out on April 26, 2020, at the Karolinska University Hospital, Stockholm, Sweden.

Nasopharyngeal swabs were collected from 23 patients suspected to have COVID-19. In 17 of these, an RT-PCR assay for SARS-CoV-2 (Corman et al., 2020) yielded a positive result. Viral RNA was extracted and cDNA was synthesized using the QIAseq FX Single Cell RNA Library Kit (QIAGEN). Illumina libraries with 350 bp fragments for shotgun sequencing were prepared using the ThruPLEX DNA-seq kit (Rubicon Genomics) and sequenced using the Illumina MiSeq technology (2×300bp).

Low quality sequences and adapters were removed using Trim Galore (version 0.4.1), followed by the removal of human sequences using Bowtie (version 2.3.2) (Langmead and Salzberg, 2012) and the human genome reference GRCh38.p13 (GenBank accession number GCA_000001405.28). The remaining reads were mapped to the complete genome of SARS-CoV-2 Wuhan-Hu-1 (GenBank accession number MN908947.3) using Bowtie. The mapped reads were subsequently assembled using the Genome Detective Virus Tool (version 1.126) (Vilsker et al., 2019) using SARS-CoV-2 (GenBank accession number MN908947.3) as reference (Table 1) and nucleotide variants were assigned for each genome (Table 2).

**Table 1.** Detailed information about the assemblies of SARS-CoV-2 strains identified in Sweden.

| Assignment | Contigs | Reads | Coverage (%) | Depth of Coverage | Identity (%) |  | Assembly length |
| --- | --- | --- | --- | --- | --- | --- | --- |
|  |  |  |  |  | NT | AA |  |
| P17157_1021 | 1 | 332800 | 99.8 | 1718.2 | 99.96 | 99.93 | 29829 |
| P17157_1007 | 1 | 210936 | 99.8 | 1180.9 | 99.95 | 99.91 | 29829 |
| P17157_1020 | 1 | 176513 | 99.7 | 935.5 | 99.96 | 99.93 | 29811 |
| P17157_1016 | 1 | 151409 | 99.7 | 776.8 | 99.97 | 99.95 | 29822 |
| P17157_1018 | 17 | 1949 | 80.6 | 12.2 | 99.73 | 99.44 | 24099 |
| P17157_1004 | 6 | 30 | 6.9 | 2.2 | 99.42 | 98.61 | 2078 |
| P17157_1005 | 5 | 30 | 6.7 | 2.5 | 99.85 | 99.61 | 1990 |
| P17157_1023 | 4 | 19 | 3.2 | 3.1 | 99.38 | 98.47 | 969 |
| P17157_1008 | 1 | 6 | 1.4 | 2.4 | 100.00 | 100.00 | 411 |

**Table 2.** List of mutations detected in the SARS-CoV-2 strains in this study.

| Nucleotide<br>Position | Ref.<br>Base | Mutant<br>Base | P17157<br>_1021 | P17157<br>_1020 | P17157<br>_1016 | P17157<br>_1007 | P17157<br>_1018* | P17157<br>_1004* | Gene | Mutation |
| --- | --- | --- | --- | --- | --- | --- | --- | --- | --- | --- |
| 219 | G | T | Yes | - | - | - | - | - |  | Noncoding |
| 241 | C | T | Yes | Yes | Yes | Yes | Yes | - |  | Noncoding |
| 1059 | C | T | Yes | Yes | Yes | - | Yes | - | ORF1a | T265I |
| 2659 | G | T | - | - | - | Yes | - | - | ORF1a | K798N |
| 2755 | G | T | - | Yes | - | - | - | - | ORF1a | Synonymous |
| 3037 | C | T | Yes | Yes | Yes | Yes | Yes | - | ORF1a | Synonymous |
| 3184 | A | G | - | - | Yes | - | - | - | ORF1a | Synonymous |
| 5147 | C | T | - | - | - | Yes | - | - | ORF1a | R1628C |
| 6285 | C | T | - | - | - | Yes | - | - | ORF1a | T2007I |
| 6352 | G | T | - | Yes | - | - | - | - | ORF1a | K2029N |
| 9193 | A | G | - | - | - | Yes | - | - | ORF1a | Synonymous |
| 11083 | G | T | - | - | - | Yes | - | - | ORF1a | L3606F |
| 11398 | T | G | - | - | - | Yes | - | - | ORF1a | Synonymous |
| 12915 | C | T | Yes | - | - | - | - | - | ORF1a | T4217I |
| 14408 | C | T | Yes | Yes | Yes | Yes | Yes | - | ORF1ab | P4715L |
| 21575 | C | T | - | - | Yes | - | - | - | S | L5F |
| 23202 | C | T | Yes | - | - | - | - | - | S | T547I |
| 23403 | A | G | Yes | Yes | Yes | Yes | Yes | - | S | D614G |
| 23463 | G | A | - | Yes | - | - | - | - | S | R634H |
| 24368 | G | T | Yes | Yes | Yes | - | - | - | S | D936Y |
| 25563 | G | T | Yes | Yes | Yes | - | Yes | - | ORF3a | Q57H |
| 27549 | C | T | - | - | - | Yes | - | - | ORF7a | Synonymous |
| 28881 | G | A | - | - | - | Yes | - | - | N | R203K |
| 28882 | G | A | - | - | - | Yes | - | - | N | R203K |
| 28883 | G | C | - | - | - | Yes | - | - | N | G204R |
| 28889 | T | C | - | Yes | - | - | - | Yes | N | S206P |
| 29287 | A | T | Yes | - | - | - | - | - | N | K338N |

\*Only accepted mutations found in the complete genomes.

SARS-CoV-2 reads were detected in nine of the samples, with variable coverage. Near complete genomes could be assembled from four samples, with a median of 29825.5±7.4 bp in length, covering 99.7-99.8% of the reference genome and 100% of the coding region, with a depth of coverage ranging from 776.8 to 1718.2. An additional genome assembly covered 80.6% of the reference genome, in 17 contigs with an average of 12.2 (Table 1).

Compared with the reference strain (GenBank accession number MN908947.3), the four complete genomes each have between 9 to 14 single nucleotide differences. All four genomes have mutations in noncoding position 241 C to T, and three mutations in coding regions, two C to T in positions 3037 and 14408, and one A to G in the position 23403 (Table 2).

The variant at 23463 bp, found in patient P17157_1020, was not found in any other SARS-CoV-2 genome that was present in GISAID and Genbank at the time that this report was drafted (13 July 2020). The impact of this spike protein R364H variant (Figure 1) was predicted by the DUET Web server (Pires et al., 2014) to have destabilizing effects. The variant is located at the surface of the S1 subunit, and could possibly affect the attachment of the virion to the cell, even though it does not change the receptor-binding domain itself. We compared the four complete SARS-CoV-2 genomes from Stockholm with genomes detected globally using the three main methods available to classify relationships between different genetic variants of the virus: GISAID, Nextstrain, and PANGOLIN (Hadfield et al., 2018; Rambaut et al., 2020). The Stockholm genomes belonged to two genetic groups: 20C/B.1/G and 20B/B.1.1/GR (Table 3). These groups were reported by the Public Health Agency of Sweden as two of the three main genetic groups found in Sweden (FHM, 2020). Somewhat surprisingly, three of the completed genomes described here are from group B.1/G, which was reported to have declined in prevalence by late April. This lineage has spread to more than 20 countries in Europe, the Americas, Asia, and Australia and corresponds to the Italian outbreak.

**Figure 1.**
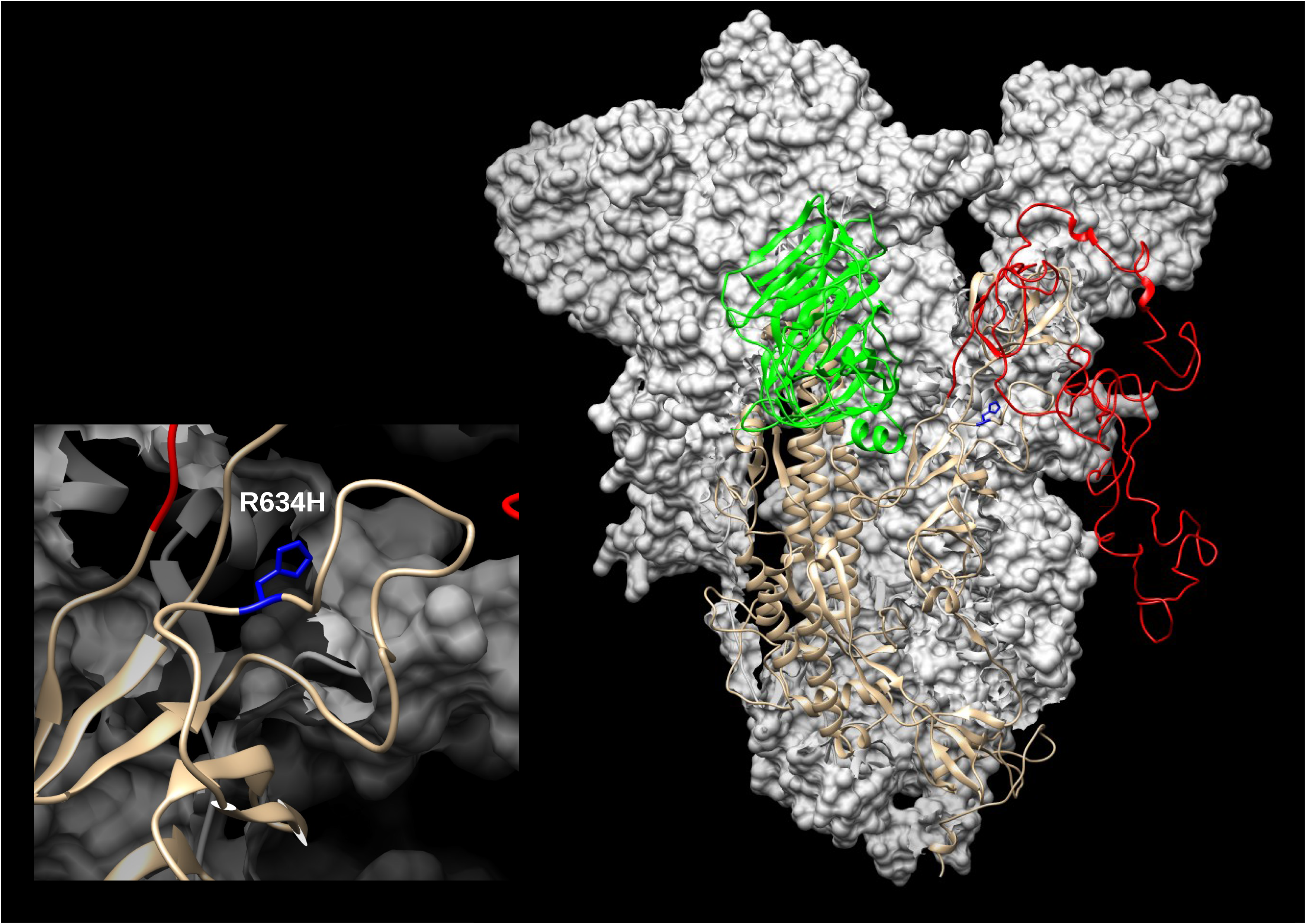
SARS-CoV-2 spike (S) modelled with SWISS-MODEL (Waterhouse et al., 2018) using 6ZGH structure as a template, drawn and colored in UCSF Chimera (Pettersen et al., 2004). N-terminal domain (NTD) is colored green and Receptor-binding domain/C-terminal domain (RBD/CTD) is red. The enlarged inset shows the location of R634H mutation (blue).

## Data availability

These sequences have been deposited in European Nucleotide Archive (ENA) under the study reference number PRJEB39632.

